# Bigger Is Fitter? Quantitative Genetic Decomposition of Selection Reveals an Adaptive Evolutionary Decline of Body Mass in a Wild Rodent Population

**DOI:** 10.1101/038604

**Authors:** Bonnet Timothée, Wandeler Peter, Camenisch Glauco, Postma Erik

**Affiliations:** Department of Evolutionary Biology and Environmental Studies, University of Zurich, Zurich, Switzerland; Natural History Museum Fribourg, Fribourg, Switzerland; Centre for Ecology and Conservation, College of Life and Environmental Sciences, University of Exeter, Cornwall Campus, Penryn, United Kingdom

## Abstract

In natural populations, quantitative trait dynamics often do not appear to follow evolutionary predictions: Despite abundant examples of natural selection acting on heritable traits, conclusive evidence for contemporary adaptive evolution remains rare for wild vertebrate populations, and phenotypic stasis seems to be the norm. This so-called ‘stasis paradox’ highlights our inability to predict evolutionary change, which is especially concerning within the context of rapid anthropogenic environmental change. While the causes underlying the stasis paradox are hotly debated, comprehensive attempts aiming at a resolution are lacking. Here we apply a quantitative genetic framework to individual-based long-term data for a wild rodent population and show that despite a positive association between body mass and fitness, there has been a genetic change towards lower body mass. The latter represents an adaptive response to viability selection favouring juveniles growing up to become relatively small adults, i.e. with a low *potential* adult mass, which presumably complete their development earlier. This selection is particularly strong towards the end of the snow-free season, and it has intensified in recent years, coinciding which a change in snowfall patterns. Importantly, neither the negative evolutionary change, nor the selective pressures that drive it, are apparent on the phenotypic level, where they are masked by phenotypic plasticity and a non-causal (i.e. non-genetic) positive association between body mass and fitness, respectively. Estimating selection at the genetic level thereby enabled us to uncover adaptive evolution in action, and to identify the corresponding phenotypic selective pressure. We thereby demonstrate that natural populations can show a rapid and adaptive evolutionary response to a novel selective pressure, and that explicitly (quantitative) genetic models are able to provide us with an understanding of the causes and consequences of selection that is superior to purely phenotypic estimates of selection and evolutionary change.

## Introduction

Given the rapid anthropogenic environmental changes experienced by organisms around the world, there is an increasing need for an ability to understand and predict the evolutionary dynamics of wild populations [1, 2]. Despite good empirical examples of the adaptive evolution of traits with a simple genetic architecture [3–5], the picture is very different for quantitative traits, which in most cases are a function of many genes of small effect [6]. Although these are the traits that are of most interest to evolutionary biologists [7, 8], predictive models of quantitative trait evolution have largely failed when applied to data from wild populations [9].

Although there is an abundant literature showing that both directional selection [10, 11] and heritable genetic variation [12, 13] are common, these pre-requisites of Darwinian evolution rarely allow us to explain evolutionary trends retrospectively, let alone to make predictions for the future [9, 14]. For example, both natural and sexual selection almost universally favour larger and heavier individuals [15]. Furthermore, size-related traits are generally moderately heritable [12, 13], and averaged across the 151 estimates compiled in [13], the heritability of body mass is 0.33 ± 0.02. These levels of directional selection and heritability generate an expectation of rapid evolution of larger body size. Nevertheless, while species do tend to get larger over geological timescales [16–19], this rate of evolution is orders of magnitude slower than predicted from the strength of selection and heritability observed in contemporary populations [9, 20, 21].

On the whole, conclusive evidence for the contemporary adaptive evolution of quantitative traits in wild vertebrate populations is remarkably scarce and elusive [9, 14], and good examples –Trinidadian guppy life-histories [22], human reproductive timing and educational attainment [23, 24], timing of pink salmon migration [25] and big-horn sheep horn size [26]—can be counted on the fingers of one hand. Furthermore, of these studies, those dealing with humans might not be representative of wild populations, [26] reported a response to harvesting-induced, artificial rather than natural selection, and despite considerable effort to uncover any evolutionary consequences of climate change [2, 27–29], only [25] were able to demonstrate an adaptation to climate.

Our apparent inability to reconcile predictions of evolutionary change based on estimates of selection and genetic variation with the (lack of) of phenotypic change observed, i.e. the ‘stasis paradox’ [9], is a major concern in urgent need of a resolution. Given how commonly evolutionary predictions fail to capture observed trait dynamics, some have concluded that there are fundamental problems that prohibit the application of quantitative genetic methods to natural populations [30, 31]. However, there are in fact three theoretically well-developed (quantitative genetic) explanations for this mismatch.

First, many studies base their conclusion of evolutionary stasis on the observed phenotypic trend not being significantly different from zero, but they do not explicitly compare the observed change to the change predicted [32]. Hence, at least some cases of phenotypic stasis may be the result of a lack of statistical power. Additionally, evolutionary change does not need to be apparent at the phenotypic level. Instead, it may be masked by phenotypically plastic changes, which may be several-fold larger and/or go opposite to the genetic change [33]. For instance, while a change in the environment may generate selection favouring an increase in the frequency of alleles promoting fat accumulation, at the same time this may create a food shortage, leading to a plastic decrease of fat reserves. As a consequence, an evolutionary trend may be masked by a counteracting plastic change, resulting in “cryptic evolution” [34, 35].

Second, the observed rates of phenotypic change (which as outlined above may provide poor estimates of evolutionary change) are typically compared to predictions from the univariate breeder’s equation, i.e. the product of selection and heritability, where selection is quantified as the phenotypic covariance between the trait of interest and fitness. In natural systems, selection however rarely acts on traits in isolation [36]. If these traits are genetically correlated to the focal trait, they may significantly alter the focal trait’s evolutionary trajectory [37, 38]. While the role of genetic correlations among traits within the same individual [39] or between the sexes [40] has received substantial attention, the potential role of genetic constraints resulting from genetic correlations between traits expressed in different life-stages has received far less attention. In particular parent-offspring conflict, e.g. a genetic trade-off between offspring quality and fecundity [41], may constrain the evolution of size [42], with positive directional selection on offspring size counterbalancing selection against investment per offspring on the level of parents [43].

Finally, even in the absence of selection on correlated traits, it is challenging to obtain an estimate of the strength of natural selection that is unbiased by the existence of a third, non-genetic variable that influences both the trait and fitness [44]. Although the univariate breeder’s equation assumes that the covariance between phenotype and fitness is solely the result of a causal relationship between the two, this assumption is likely to be violated, especially in natural populations [14, 45]. For instance, a trait that plastically responds to food availability, such as body mass, is likely to covary with fitness at the phenotypic level, irrespective of the causal effects of this trait on fitness: individuals that have access to more food are heavier and reproduce more [37, 46].

However, because the fitter individuals are not genetically different in terms of body mass, this covariation has no evolutionary consequences, even if body mass is heritable [44].

While these difficulties have been discussed previously, and studies regularly note that the mismatch between the observed and predicted response may be attributable to any of them, they are rarely accounted for in an explicit, quantitative manner. Therefore, we here apply a comprehensive analytical framework to long-term individual-based body mass data for a wild rodent population, which appear to show a mismatch between the observed rate of phenotypic change and the predicted rate of genetic change. To resolve this mismatch, we use information on within-population relatedness and individual-level trait measurements [47, 48] to obtain a statistically robust estimate of the direction and rate of genetic change [14, 49, 50]. Subsequently we disentangle the role of genes and the environment in shaping the covariance between body mass and fitness, and identify the likely target of selection. This allows us to directly compare the observed genetic change to a range of evolutionary predictions, and to thereby resolve the stasis paradox and provide a deeper understanding of selection and evolution in this system.

## Results and discussion

Based on ten years of data on an alpine population of snow voles [51, 52] (*Chionomys nivalis*, Martin 1842), we find that relatively heavy individuals both survive better (*p* = 0.04) and produce more offspring per year (*p* = 0.003). Assuming causality, this generates strong phenotypic selection favouring heavier individuals (selection differential *S* = 0.86 g, *p* < 10^−5^; mean adult mass= 41.7 g, standard deviation=5.2 g). In line with other morphological traits [12, 13], variation in body mass has a significant additive genetic component (*V_A_*=4.34 g^2^, 95%CI [2.40;7.36]), which corresponds to a heritability (*h*^2^) of 0.21 (95%CI [0.11;0.29]). Similarly, we find significant additive genetic variance in fitness (*V_A_*=0.10; 95%CI [0.06;0.19], *h*^2^ = 0.06 95%CI [0.04;0.12]), measured as relative lifetime reproductive success (rLRS).

Given these estimates of selection (*S*) and heritability (*h*^2^), and after normalizing by a mean generation time of 1.2 years, the breeder’s equation (*R* = *h*^2^*S*) predicts an adaptive evolutionary response (*R*) in body mass [14, 48], i.e. an increase in the mean breeding value for body mass over time, of 0.17 g/year (95%CI [0.07;0.28]; Fig 2A UBE, Univariate Breeder’s Equation). However, over the past nine years (approximately eight generations), the change in mean body mass is not significant and small at best (0.08 g/y; 95%CI [-0.02;0.18]; p=0.14) after correcting for changes in demographic structure (i.e. accounting for sex and age effects, see Fig 1A and S1). This apparent mismatch between the predicted increase in body mass based on the breeder’s equation and the absence of a strong and statistically significant phenotypic change would appear to provide yet another example of the stasis paradox [9].

**Figure 1.**
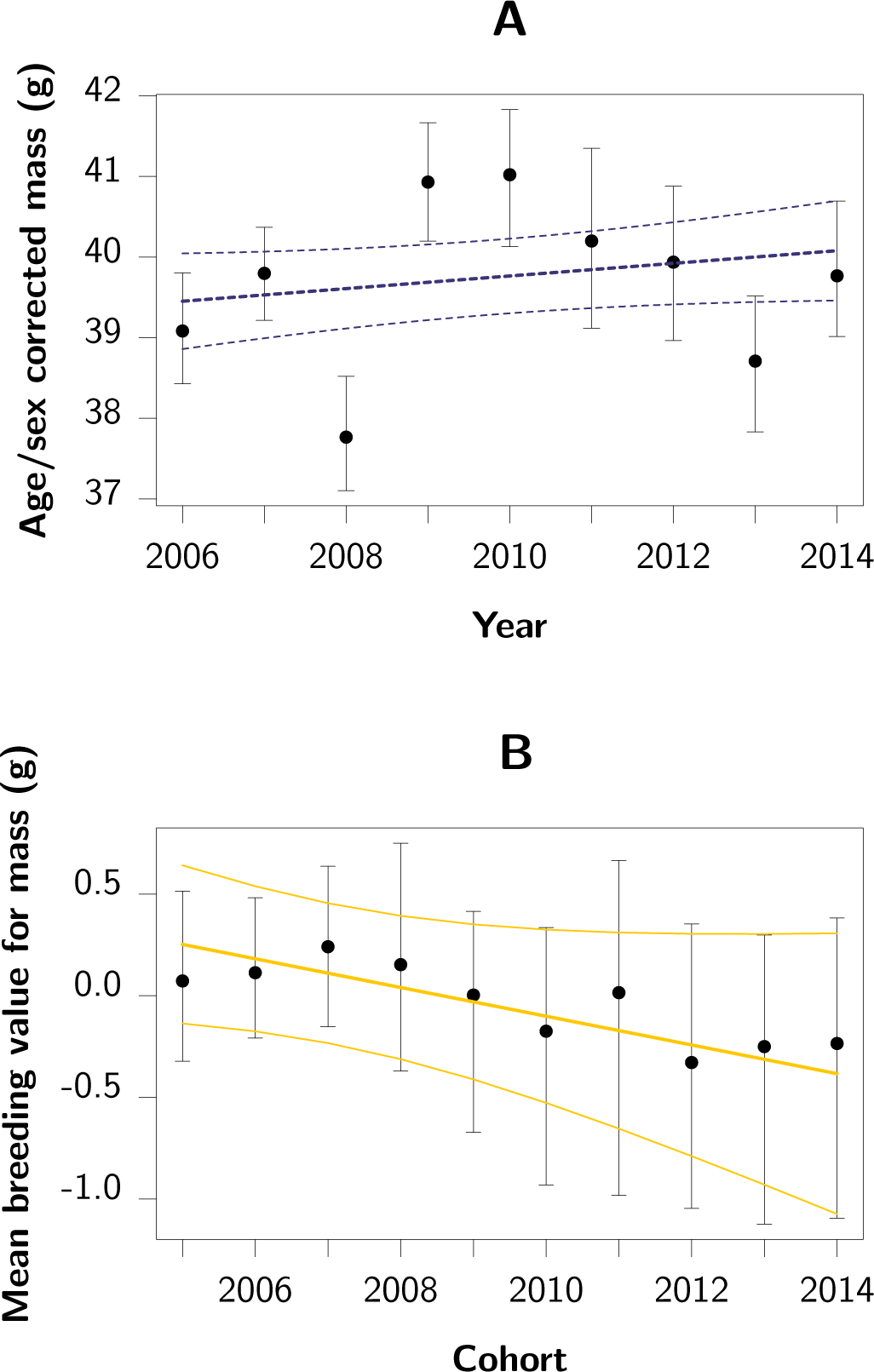
Temporal variation in mass and estimated breeding values for mass. (**A**): Year-specific mean mass corrected for age, sex and date of measurement, with 95%CI. (**B**): Cohort-specific mean estimated breeding value for mass with their 95%CI and the trend in breeding value with 95%CI. Note the different scaling of the y-axes in **A** and **B**.

To test whether the predicted positive genetic trend, i.e. an increase in breeding values, is being masked by the effects of phenotypic plasticity [9, 35], we directly estimated the additive genetic covariance between mass and fitness. Based on the Robertson-Price’s equation, this provides an unbiased estimate of the rate of genetic change per generation [14, 53, 54]. Contrary to the predicted evolutionary increase in body mass, this estimate of genetic change in mass is strongly negative and highly significant (*p*_MCMC_ < 0.001; Fig 2A GCPE, Genetic Change Price Equation). When normalized by a mean generation time of 1.2 years, this provides a rate of evolutionary change of −0.29 g/year (95%CI [−0.55; −0.07]) or approximately 7,300 Darwins (the natural logarithm of the proportional change in trait value over a million years [55]), which is in line with other estimated rates of “micro-evolution” (e.g. between 3,700 and 45,000 Darwins in the Trinidadian guppies [56]). Importantly, this rate of evolution is unlikely to have happened solely through genetic drift (*p*_MCMC_ < 0.001; Fig S2 and S3) [50], and therefore most likely reflects a response to selection favouring genetically lighter individuals.

**Figure 2.**
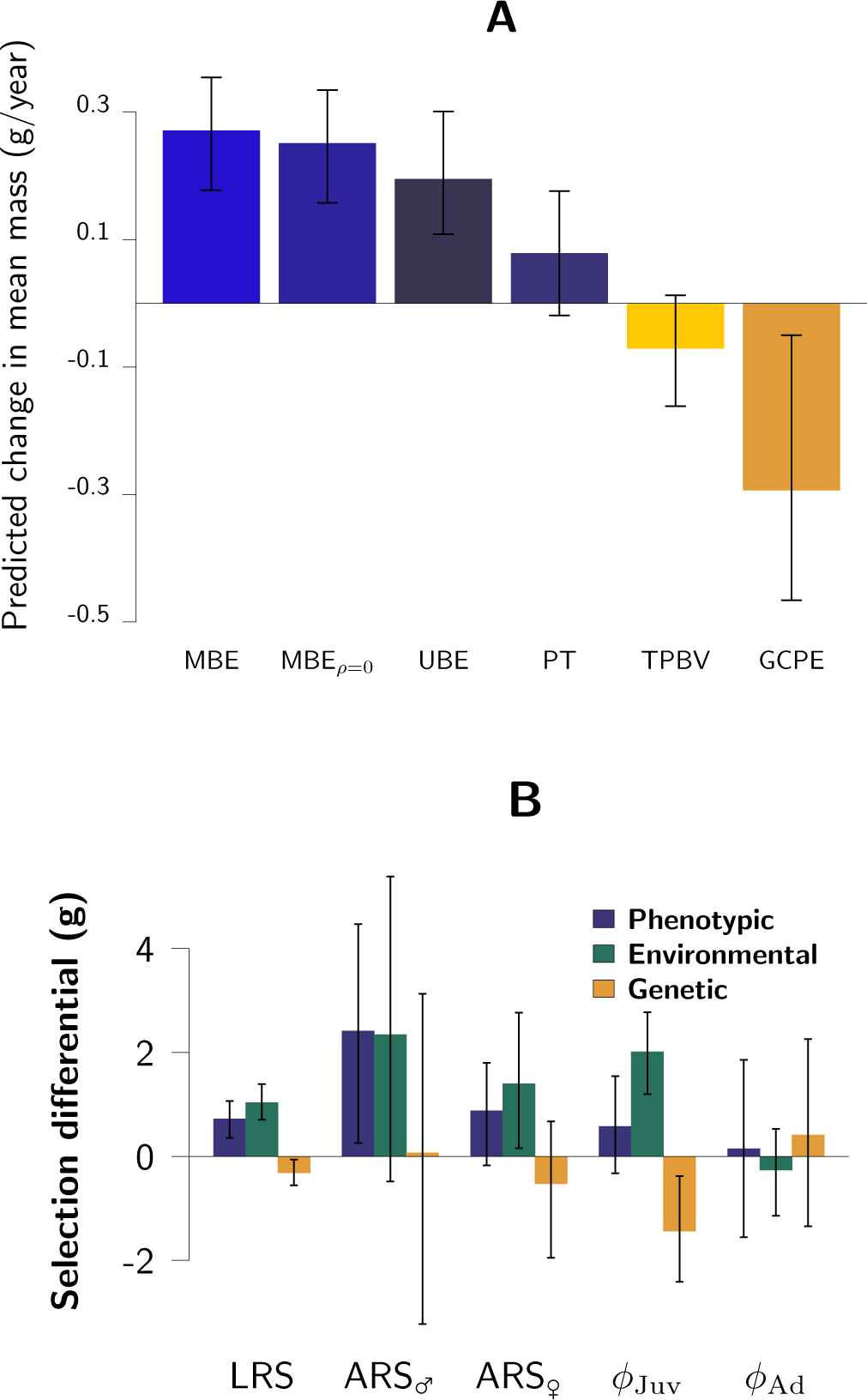
Predicted and observed rates of evolutionary change. (**A**): Rates of evolutionary change predicted by (from left to right) the breeder’s equation in its multivariate form (MBE), the multivariate breeder’s equation while constraining the genetic correlations to zero (MBE_*ρ*=0_), and the univariate breeder’s equation (UBE), followed by the phenotypic trend (PT), the trend in predicted breeding values (TPBV) and the genetic change estimated by the Price equation (GCPE). (**B**): Phenotypic, genetic and environmental selection differential for total selection (LRS), fertility selection in males (ARS_♂_) and females (ARS_♀_), viability selection in juveniles (*ϕ*_Juv_) and in adults (*ϕ*_A_). Both panels show posterior modes, with vertical lines indicating 95%CI.

The above result was confirmed by an independent estimate of the rate of evolution using best linear unbiased predictors (BLUPs) of breeding values for mass: Taking into account the non-independence of BLUPs and sampling variance [49, 50], we find that predicted breeding values have likely declined over the past nine years (−0.07 g/year, 95%CI [−0.16; 0.01], *p*_MCMC_=0.06; Fig 1B & Fig 2A TPBV, Trend in Predicted Breeding Values), and this despite the BLUPs approach being potentially biased towards the phenotypic trend [49] (i.e. in this case towards zero). This negative trend, combined with the fact that the phenotypic mean has either remained constant or has shown a slight increase (see above), implies that the plastic component of body mass must have increased. Although the cause of this increase remains unknown, population size has declined over the study period (Fig S1), which may have resulted in an increase in the per-capita resource availability (i.e. density dependence). Alternatively, the absolute food availability or quality may have improved. Interestingly, although these environmental changes may be coincidental, they may also be a direct result of a change in the selection regime or the evolutionary change toward smaller size [35, 57].

As the phenotypic selection differential (*σ_P_*(*m*, *ω*), i.e. the phenotypic covariance between body mass *m* and fitness *ω*, measured as relative lifetime reproductive success (rLRS)) is equal to the sum of the additive genetic and environmental covariances between mass and fitness (*σ_A_*(*m*, *ω*) and *σ_E_*(*m*, *ω*), respectively [14, 53, 54, 58, 59], it follows that because *σ_P_*(*m*, *ω*) is positive and *σ_A_*(*m*, *ω*) is negative, the environmental covariance must be large and positive (Fig 2B LRS). In other words, while environmental conditions that make voles heavy (for instance abundance of food or lack of parasites) also make them successful at reproducing and surviving, there is no causal *positive* relationship between breeding values for mass and fitness. It is this difference in sign between *σ_A_*(*m*, *ω*) and *σ_E_*(*m*, *ω*) that represents an extreme violation of the assumption of the breeder’s equation that *σ_A_*(*m*, *ω*)/*V_A_* = *σ_E_*(*m*, *ω*)/*V_E_* [44]. Hence, our initial prediction of evolution based on phenotypic estimates of selection was wrong, illustrating how these may provide severely biased predictions of the evolutionary trajectories of wild populations. But *why* is evolution taking place in a direction that is opposite to apparent phenotypic selection?

Indirect selection may be acting on body mass through one or more traits genetically correlated to mass [36, 38]: a positive selective pressure on a negatively correlated trait (or a negative selective pressure on a positively correlated trait) would indirectly select for lower mass. However, the genetic correlations among the three morphological traits for which we have data—body mass (*m*), body length (*b*) and tail length (*t*)—are all positive (estimates and 95%CI: *ρ_m,b_* = 0.79 [0.06; 0.93]; *ρ_m,t_* = 0.40 [0.01; 0.66]; *ρ_t,b_* = 0.56 [−0.04; 0.85]), and direct selection is positive for two of them and only slightly negative for one of them (see table S1). Hence, the predicted response based on the multivariate breeder’s equation (Fig 2A MBE, Multivariate Breeder Equation) is very similar to that based on its univariate counterpart (Fig 2A UBE), as well as to that based on a multivariate breeder’s equation constraining the correlations to zero (Fig 2A MBE_*ρ*=0_). Therefore, the genetic correlations among the traits considered do not constrain the evolution of body mass and do not explain the trend towards lower body mass. To test whether parent-offspring conflict between size and fertility might be constraining the evolution of size, we furthermore estimated the genetic correlation between juvenile size and adult annual reproductive success, which should be negative for it to provide a constraint [43]. Although we were not able to incorporate all sources of uncertainty into the estimation of this correlation (see Methods), our best estimate was 0.21, 95%CI [−0.24; 0.74]), which argues against a major role for a trade-off between fertility and offspring size in driving the observed evolution toward smaller sizes.

To pursue the possibility that the counter-intuitive direction of evolution was due to selective pressures directly acting on body mass, we first tested which fitness component is negatively associated with genes for being heavy. Computing sex- and age-specific genetic covariances between mass and fitness components revealed that whereas the genetic covariances between mass and both relative annual reproductive success and adult survival are close to zero in both sexes (Fig 2B), the genetic covariance between mass and over-winter survival is negative in juveniles (−0.98 [-2.44;−0.18] on a logit scale, *p*_MCMC_=0.01). Because the genetic correlation between juvenile and adult mass is positive (*r_A_* = 0.88; 95%CI [0.39;1]) and significantly different from 0 (p=0.004) but not 1 (p=0.35), selection on juvenile mass can shape genetic variance for mass at all ages, and thereby contribute to the observed negative genetic change [60]. While this shows that negative viability selection of juvenile mass is responsible for the genetic change toward smaller individuals, how come survival is higher for heavier phenotypes *and* lighter genotypes?

Juvenile mass covaries positively with both within- and between-year survival (*p* = 0.009 and *p* = 1.3 × 10^−6^, respectively). However, juveniles can only be captured when they first leave their burrow, at an age of approximately three weeks [61], when they weigh 12 to 20 g. They may however continue to be captured until the end of the season, when they can reach weights of up to 50 g. Because of growth, mass measurements are therefore not directly comparable among juveniles of different ages. Indeed, at least part of the positive phenotypic relationship between juvenile mass and survival probability is likely to be mediated by the fact that both increase with age [62]. In addition, viability selection introduces non-random missing data, which further biases estimates of viability selection on mass [62, 63]. Therefore, the positive phenotypic association between juvenile body mass and survival is unlikely to be the result of an among-individual covariation of survival probability and mass, and hence provides a poor representation of selection.

The (co)variance decompositions presented above have the advantage that they do not make causative statements. For example, a genetic covariance describing the rate of evolution has a self-contained, tautological, meaning and does not make any assumptions with respect to its causes [64]. However, if we are to identify the target of juvenile viability selection, we must adopt a more traditional hypothesis testing framework. This is somewhat complicated by our relatively poor understanding of snow vole biology, much of which takes place below the rocks or the snow, and the impossibility to carry out manipulative experiments in a natural setting. Although this means that the scenario detailed below is in part speculative, it is well-supported by our data.

We hypothesised that juveniles with a low *potential adult* mass, i.e. juveniles that will grow up to be relatively small adults (*if* they survive), require less time to reach their adult size. Furthermore, when the period favourable for growth is limited, and juveniles that have not completed development before the arrival of winter pay a survival cost, for example due to trade-offs between growth and vital physiological processes [65, 66], survival selection will favour juveniles that need less time to reach their adult size. This scenario generates selection for small size, especially for juveniles born toward the end of the season (Fig 3A and Fig S4), and gives rise to a negative genetic association between mass and survival. On the phenotypic level this negative selection would however be masked by the positive within-individual age-related association between mass and survival (see above).

**Figure 3.**
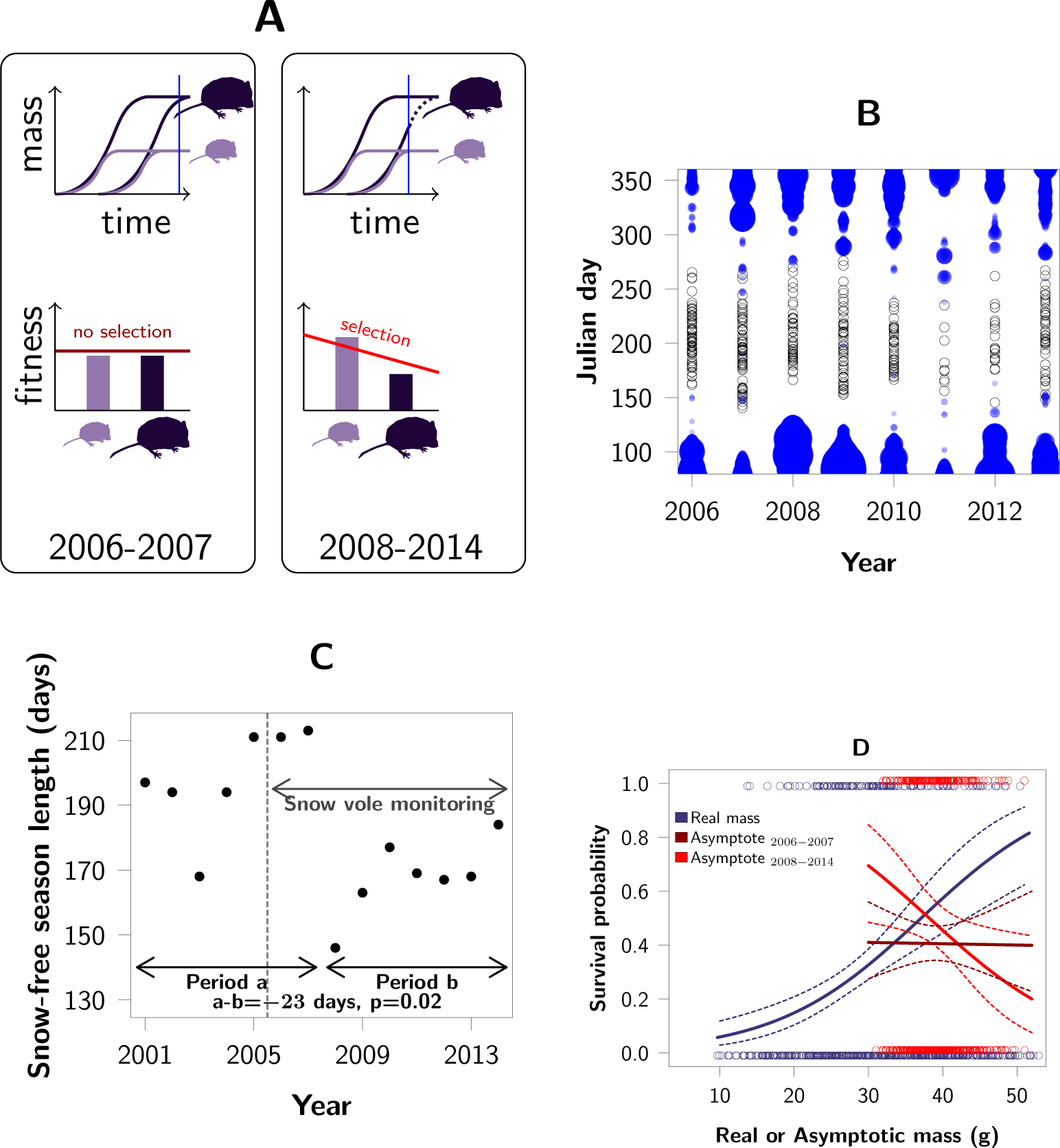
Snow-free season, timing of reproduction and selection for potential adult mass. (**A**) Hypothetical scenario generating selection for lower body mass: In years with short summers (2008-2014), juveniles born late and having a large potential adult mass are still growing at the onset of winter and therefore less likely to survive (blue vertical line). This results in selection for individuals with a low potential adult mass, despite mass covarying positively with survival on a within-individual level due to variation in age. (**B**) Births (black dots) only occur during the snow free season (the depth of the snow cover is indicated by the thickness of the blue dots), (**C**): which in 2008-2014 has been shorter than in the preceding 7 year. Therefore, (**D**) despite a positive phenotypic selection on body mass (blue), predicted adult mass was selectively neutral in 2006-2007 (brown), and was negatively selected in 2008-2014 (red).

Although, as we emphasised above, inferring a causal relationship between a trait and fitness based on their covariance requires great care, we set out to obtain an estimate of viability selection that is unbiased by growth and non-random missing data due to mass-dependent mortality occurring after the first capture [62]. To this end we used a Bayesian model to simultaneously infer birth dates and growth curves for all juveniles observed at least once. Although we cannot account for viability selection acting before the first capture, this model enabled us to quantify viability selection on age-corrected juvenile mass—i.e. asymptotic or predicted adult mass—, and thereby compare all individuals at the same developmental stage (adulthood), irrespective of their fate. Inferred birth dates revealed that snow fallen during the preceding winter is a major ecological factor constraining the onset of reproduction in spring, with reproduction starting on average 40 days after the snow has melted (SE 4.5, *p* = 4 × 10^−5^) (Fig 3B). As a consequence, juveniles only have a limited amount of time to grow and reach their adult mass before the return of winter. As growth rate and predicted adult mass are only weakly (and negatively) correlated (correlation −0.077; 95%CI [−0.150; −0.002]), juveniles with a smaller adult mass on average require less time to complete development. The strength of survival selection acting on predicted adult mass was slightly negative when averaged over all years and the complete reproductive season (*p*_MCMC_=0.13), but interacted strongly and significantly with the number of days between birth and the first snowfall of that year (interaction = 0.0025, 95%CI [0.0001; 0.0048] *p*_MCMC_=0.008; for other effects see table S2). This implies that individuals born closer to the first snowfall are more strongly selected for a low adult mass, and that the length of the snow-free period in a given year determines the total selection experienced by the population in that year.

At our field site, the length of the snow-free period in the years 2008 to 2014 has been significantly shorter than during the preceding seven years (Fig 3C). The latter coincides with a period of exceptionally high snowfall, low temperatures and a long duration of snow cover, across the Swiss Alps [67]. Our model estimates that in 2006 and 2007, when the snow-free period was long, most juveniles reached their adult mass before the first snowfall, and there was hence no selection on predicted adult mass (*β* = −0.002, SE= 0.0006, *p*_MCMC_=0.47, Fig 3D; D; S4). However, in all subsequent years, the snow-free period was much shorter, and there was selection for a lower predicted adult mass (*β* = −0.10, SE=0.0008, *p*_MCMC_=0.009). This suggests that the shortening of the snow-free season, and thereby selection for lower predicted adult mass, is a novel phenomenon that the population is currently in the process of adapting to. Assuming that the additive genetic variation in potential adult mass is similar to the additive genetic variation in mass (4.34 g^2^, see above), the breeder’s equation (*R* = *βV_A_*) predicts a response to selection on potential adult mass between −0.72 and −0.24 g/year (from 2008 and 2014) which is in line with the observed rate of body mass evolution (−0.29 g/year [−0.55; −0.07] from 2006 to 2014).

Model complexity and data availability prohibit rigorously disentangling genetic and environmental sources of variation in predicted adult mass among individuals and over time. Therefore we cannot rule out the possibility that the selective pressure we identified is not causative [68], nor can we rule out other selective forces contributing to the observed evolution. Nevertheless, the hypothesis presented here is consistent with observations: The cohort born in 2013 had an estimated adult mass that was 1.02 g smaller than the cohort born in 2006 (p=0.05). This decline in mass is predicted to have increased population-level juvenile survival by 2.5%, and may have contributed positively to population recovery (Fig S1).

## Conclusion

We have exploited a case of apparent evolutionary stasis to gain a deeper insight into the evolutionary dynamics of natural populations, and the selective pressures that shape them. Whereas estimates of selection and evolution based on phenotypic data alone can easily mislead our understanding of the selective and evolutionary processes in natural populations, a quantitative genetic framework applied to individual-based long-term data allows us to unravel evolutionary and environmental changes over time, and to obtain unbiased estimates of selection. This has resolved a case of apparent evolutionary stasis, providing a comprehensive empirical demonstration of contemporary adaptive evolution in response to a climatic fluctuation.

## Methods

### Snow vole monitoring

Monitoring of the snow vole population began in 2006 and the present work uses data collected until the fall of 2014. The snow vole monitoring was authorised by the *Amt für Lebensmittelsicherheit und Tiergesundheit*, Chur, Switzerland. The study site is located at around 2030m above sea level, in the central eastern Alps near Churwalden, Switzerland (46°48’ N, 9°34’E). The site consists of scree, interspersed with patches of alpine meadows and surrounded by habitat unsuitable for snow voles: a spruce forest to the West, a cliff to the East and large meadows to the North and South. In accordance with it being considered a rock-dwelling specialist [61], at our study site snow voles were almost never captured outside of the rocky area. Given that it is ecologically fairly isolated, we were able to monitor the whole population. Trapping throughout the whole study area took place during the snow-free period, between late May and mid-October. One trapping session consisted of four trapping nights. Analyses presented here are based on a total of two (in one year), three (in three years) or five (in five years) trapping sessions per season. All newly-captured individuals weighing more than 14 g were marked with a subcutaneous passive transponder (PIT, ISO transponder, Tierchip Dasmann, Tecklenburg). Additionally, an ear tissue sample was taken (maximum 2 mm in diameter) using a thumb type punch (Harvard Apparatus) and stored in 100% ethanol at −20°C. DNA extracted from these samples was genotyped for 18 autosomal microsatellites developed for this population [69], as well as for the *Sry* locus to confirm the sex of all individuals. Finally, another Y-linked marker as well as a mitochondrial marker was used check for errors in the inferred pedigree (see below). An identity analysis in CERVUS v.3.0 [70] allowed us to identify animals sampled multiply, either because they lost their PIT, or because at their first capture as a juvenile they were too small to receive a PIT. Over the study period we obtained 3382 captures of 937 individuals (see Fig SI5). All the analyses were carried out in R [71]. Specific packages are referenced below.

### Pedigree inference

Parentage was inferred by simultaneously reconstructing paternity, maternity and sibship using a maximum likelihood model in MasterBayes [72]. Parentage was assigned using a parental pool of all adults present in the examined year and the previous year, assuming polygamy and a uniform genotyping error rate of 0.5% for all 18 loci. As it is known that in rare cases females reach sexual maturity in their year of birth [61], we matched the genotypes of all individuals against the genotypes that can be produced by all possible pairs of males and females. We retrieved the combinations having two or less mismatches (out of 18 loci) and ensured that parental links were not circular and were temporally consistent. This way, we identified eight young females as mothers of animals born in the same year, with a known father but a mother not previously identified. All of these females were relatively heavy (*>*33 g) at the end of the season and their home-ranges matched those of their putative offspring. Finally, the pedigree was checked using a polymorphic Y-linked locus developed for this population [73], as well as a fragment of the mitochondrial DNA control region, amplified using vole specific primers [74]. There were no inconsistencies between the transmission of these three markers and the reconstructed pedigree.

Although the study periods spans over 7.5 times the mean generation time, the final pedigree had a maximum depth of 11 generations (one for each of the nine years of monitoring, plus one for the unobserved parents of siblings that could be inferred in the first year, plus one due to the reproduction of a few females in the same year they were born) and a mean depth of 3.8 generations. The pedigree consists of 987 individuals (more than the number of captured individuals because the pedigree contained the unobserved parents of some full-siblings that could be reliably inferred), 458 full-sibling pairs, 3010 half-sibling pairs, 764 known maternities and 776 known paternities, so that, excluding the base population, 86% of the total parental links were recovered.

### Traits

The recapture probability from one trapping session to the next was estimated to be 0.924 (SE 0.012) for adults and 0.814 (SE 0.030) for juveniles using mark-recapture models. Thus, with three trapping session a year, the probability not to trap an individual present in a given year is below 10^−3^. Not surprisingly, no animal was captured in year *y*, not captured in *y* + 1, but captured or found to be a parent of a juvenile in *y* + 2 or later. Therefore, capture data almost perfectly matches over-winter survival. However, as is almost always the case in these type of studies, we are unable to separate death from permanent emigration. Importantly however, as both have the same consequences on the population level, this does not affect our evolutionary predictions.

Annual and lifetime reproductive success (ARS and LRS, respectively) were defined as the number of offspring attributed to an individual in the pedigree, either over a specific year or over its lifetime. 56 individuals born of local parents were not captured in their first year, but only as adult during the next summer, probably because they were born late in the season and we had only few opportunities to catch them. This means that we miss a fraction of the juveniles that are not observed in their first year and die, or emigrate, during the following winter. We acknowledge that this means that our measures of ARS and LRS partly conflate adult reproductive success and the viability of those juveniles that were never observed, but our measures are the most complete measures of reproductive success available in this system.

In order to estimate total selection, we used relative LRS (*ω*) as proxy for fitness [36], where 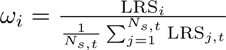. Here, *N*_*s,t*_ is the number of individuals of same sex as the focal individual *i*, present in the cohort *t*, so that 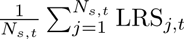 is the sex-specific, cohort-specific mean of LRS. The latter is required as the mean LRS differs between males and females due to imperfect sampling [14]. We used cohort-specific means in order to account for variations in population growth rate. In addition, to estimate the viability and fertility component of total selection, we used ARS and survival, standardized by their sex- and year-specific means.

Generation time was defined as the mean age of parents at birth of their offspring [75] and used to convert the predictions and estimations of evolution into grams per year.

Mass (*m*) was measured to the nearest gram with a spring scale. Both body length (*b*), measured from the tip of the nose to the base of the tail, and tail length, measured from the tip to the base of the tail (*c*), were measured to the closest mm with a calliper while holding the animal by the tail.

### Selection

Selection differentials were estimated using bivariate linear mixed models, as the individual-level covariance between fitness (i.e. relative LRS for total selection, relative ARS for fertility selection and relative survival for viability selection) and mass (corrected for sex, age and cohort). However, while this provides the best estimate of the within-generation change in trait mean due to selection [36], because the distribution of fitness is not Gaussian, the credibility intervals produced by these models are not exact. Hence, the statistical significance of selection was tested using a univariate over-dispersed Poisson generalized linear mixed model (GLMM) in which LRS was modelled as a function of individual standardized mass and including sex and age as covariates and cohort as a random effect. Note that the latter estimates the effect of mass on a transformed scale, and therefore cannot be directly used to quantify an effect of selection on the original scale measured in grams [76]. The significance based on the basis of the GLMM was confirmed by non-parametric bootstrapping. Similarly, we tested for the significance of selection through ARS only, using an over-dispersed Poisson GLMM including sex as a fixed effect, and year and individual as random effects.

The estimation of survival selection is facilitated by the fact that the year-to-year individual recapture probability is effectively 1. Therefore, selection on year-to-year survival was tested for by a binomial GLMM. This model included sex, age and their interaction as fixed effects, and year as a random effect.

In order to integrate the uncertainty in the estimation of selection with the uncertainty in the estimation of heritability when predicting the rate of evolution, selection differentials and gradients were also obtained from the multivariate animal model presented below.

### Quantitative genetic analyses

We used uni- and multivariate animal models to estimate additive genetic variances, covariances and breeding values [48, 77, 78] with MCMCglmm [79]. All estimations were carried out in a Bayesian framework in order to propagate uncertainty when computing composite statistics such as heritabilities and rates of genetic change [58]. All estimates provided in the text are posterior modes and credibility intervals are highest probability density intervals at the 95% level. All the animal models were run for 1,300,000 iterations with a burnin of 300,000 and a thinning of 1,000, so that the autocorrelations of each parameter chain was less than 0.1.

Convergence was checked graphically and by running each model twice.

### Univariate models

We first carried out univariate model selection, fitting models without an additive genetic effect, to determine which fixed and random effects to include. Based on AICc [80], and fitting the models by Maximum of Likelihood in lme4 [81], we obtained a model that predicts the mass *m_i,t_* of individual *i* at time *t* by: age, as a factor (juvenile or adult); sex as a factor; the interaction between age and sex; Julian dates and squared Julian dates (in order to correct for population-level seasonal variation in mass: individuals of all sexes and ages tend to be heavier in the middle of summer and lighter in spring and fall, possibly as a result of food abundance), which were centered and divided by their standard deviations in order to facilitate convergence; the interaction between age and Julian date; the interaction between sex and Julian date; the three way interaction between age, sex and Julian dates; a random intercept for individual, i.e. permanent environment; and a random intercept for year. The inclusion of year accounts for non-independence of observation within years, while the permanent environment random effect accounted for the non-independence of repeated measurements made on the same individual [82].

We then fitted an animal model by adding a random intercept modelling variance associated with mother identity [78], and a random intercept modelling additive genetic variance. Although it was not included in the best models, we kept inbreeding coefficient (estimated from the pedigree) as a covariate, because leaving it out could bias the later estimation of additive genetic variation [83]. Nevertheless, animal models fitted without this covariate gave indistinguishable estimates. The univariate animal model for body mass can be written as:

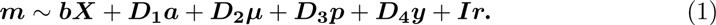

Here ***X***, ***D*_1_**, ***D*_2_**, ***D*_3_** and ***D*_4_** are design matrices relating observations to the parameters to estimate, ***b*** is a matrix of fixed effects, ***a***, ***μ***, ***p*** and ***y*** are random effects accounting for the variance associated with breeding value, mother, permanent environment (i.e. individual repeatability) and year, respectively.

The most important aspect of this model is that ***a***, the matrix of breeding values, follows a multivariate normal distribution:

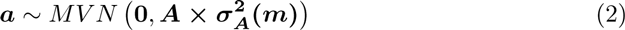

where ***A*** is the relatedness matrix describing the relatedness among all individuals, and 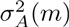 is the additive genetic variance in body mass. For univariate animal models for each random effect we used a vague proper prior with the variance parameter set to 1 and the degree of belief set to 0.002.

### Multivariate models

Univariate animal models can be expanded to multivariate models in order to estimate genetic correlations, genetic gradients and genetic differentials between body mass (*m*), body length (*l*), tail length (*t*) and fitness (*ω*).

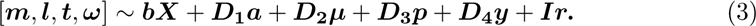

The fixed part of the model matches that used for each trait in univariate models. We used parameter-expanded priors for the variance components in order to facilitate convergence of parameters that were close to zero [79]. The working parameter prior was normally distributed, with a mean of 0 and a variance of 1000. For *ω*, the residual variance was set to zero in the model, so that the residual variance in fitness is estimated by the permanent environment random effect. Although fitness is measured only once per individual, this allows the estimation of the covariance between fitness and the non-additive genetic repeatable component of trait variation, that the environmental component of selection [14]. We assumed a Gaussian distribution for fitness so that covariances between traits and fitness could be directly interpreted as selection differentials [76].

The matrix of breeding values (*a*) here follows a multivariate normal distribution:

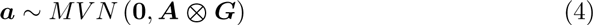

where ***A*** is the relatedness matrix describing the relatedness among all individuals, and *G* is the G-matrix, containing the additive genetic variances and covariances among all traits.

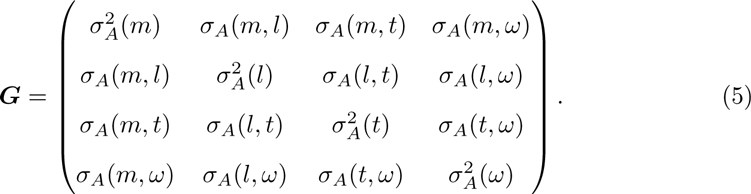

For any trait *z*, *σ_A_*(*z*, *ω*) is the genetic differential, that is, the predicted rate of evolutionary change according to Robertson’s secondary theorem of natural selection, or Price equation applied to genetic variation [14, 53, 54, 58]. The Price equation is generally presented as a prediction of evolutionary change over the next generation, but it has also been used as a description of change [64, 84, 85]. We use this prediction retrospectively, as an estimation of the mean evolutionary change that has occurred during the study period, which makes the assumption that *ω* is a good measure of fitness, because when “real fitness” is used, the equation is a mathematical tautology, i.e. it is exact [64]. A deviation from this perfect fitness measure could come from random Mendelian segregation or systematic meiosis distortion. Our results were robust to the use of an annualized measure of fitness (annual reproductive success plus twice survival), and to standardizing fitness across all individuals, within years, within cohorts, or within sexes.

For two traits *z* and *y*, the genetic correlation is 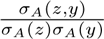. The vector of selection differentials on the three traits (***S***) was estimated as the sum of the vectors of covariances between traits and *ω* in the variance-covariance matrices for ***a***, ***p***, ***m*** and ***r***; which was equivalent to the selection differential computed in the paragraph on selection above. Let *G’* be a subset of *G* excluding the column and the row that contain *ω*. The vector of selection gradients on the three traits (*β*) was estimated as (*G′* + *P′* + *R′*)^−1^*S*, where *P′* and *R′* are the equivalent of *G′* for permanent environment effects and for residuals, respectively [38, 58]. The prediction of the multivariate breeder’s equation is obtained by 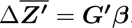. In order to visualise the effect of the genetic correlations on the predictions of evolution, we also applied a multivariate breeder’s equation that assumes the genetic correlations to be zero. To this end we multiplied the **G’** matrix by the identity matrix [38]: 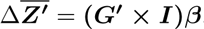. We excluded the among-year level covariance from the selection differential and gradients, because (i) covariation between mass and fitness at the level of year does not correspond to selection as it does not occur among individuals, and (ii) due to the standardization of relative fitness at the level of cohorts, the among year variance and covariances involving *ω* were effectively zero 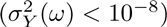.

To investigate the potential role of parent-offspring conflict, we estimated the genetic correlation between parental ARS and offspring mass using a bivariate animal model. For juvenile mass, we used predicted adult mass (i.e. age-corrected juvenile mass; see below). We were not able to incorporate the uncertainty in the estimation of predicted adult mass in this model and used the mean of the posterior distribution of each individual. The credibility intervals derived from this model are therefore only indicative. The model included sex as fixed effect, and random additive genetic, individual identify (i.e. permanent environment effects), maternal identity and year effects.

To decompose selection components into environmental and genetic covariances (as presented in Fig 2B) we fitted models like the one described in equations (1-3), but replacing fitness (*ω*) by the appropriate fitness components: relative ARS or relative survival; and taking the appropriate subset of the data: adult males (ARS_♂_), adult females (ARS_♀_), adults (*ϕ*_Ad_) or juveniles (*ϕ*_Juv_). To estimate readily interpretable environmental and genetic covariances between body mass and fitness we again assumed a Gaussian distribution of fitness components [76].

### Test of genetic correlations

We used ASReml-R [86, 87] to test the genetic correlation between mass in adults and in juveniles against 1 and 0, by considering them as two separate traits. We first ran an unconstrained model and then reran it with the genetic correlation parameter set to 0.99 (and not exactly to 1 because ASReml cannot invert matrices with perfect correlations), or 0 respectively. The fit of the unconstrained model was then compared to that of the two constrained models using a likelihood ratio test with one degree of freedom [88].

### Birth date and growth prediction

Using the Bayesian programming environment JAGS [89], we fitted a multivariate Bayesian model to mass measurements of all 613 juveniles with mass data, and to their overwinter survival. The model simultaneously estimated individual growth curves—that is onset of growth (although this is referred to as “birth date” hereafter, this actually is the projected time when mass was zero, i.e. at conception), individual growth rates and asymptotic masses (i.e. predicted adult mass) of all juveniles—and the effect of predicted adult mass on overwinter survival. The model clustered juveniles from the same mother born in the same year into litters (see e.g. [90] for a similar approach), assuming a maximum of five litters per year and assuming that successive litters are at least 20 days apart [61]. Preliminary model selection, assuming no differences in predicted adult masses among individuals, selected a monomolecular growth model (ΔDIC > 80) over Gompertz and logistic models, as defined in [91]. The model accounted for measurement error in mass, assuming that the standard deviation of the errors was that observed in animals measured multiple times on the same day (2.05g). In order to estimate the overall viability selection on predicted adult mass, we performed within the model a logistic regression of year-to-year survival on sex and predicted adult mass. In order to test for the effect of the length of snow free period on the selection on predicted adult mass, we reran the full model including time until the first snow fall and its interaction with predicted adult mass in the logistic regression. We use the estimates of these two models to predict the survival probability as a function of predicted adult mass for every year, or for groups of years, depending on the distribution of birth dates and on the timing of the first snow fall.

Three MCMC chains were run for 6,300,000 iterations, with a burnin of 300,000 and a thinning of 6,000. Convergence was assessed by visual examination of the traces, and by checking that 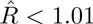 *R* < 1.01. Convergence was not achieved for the litter affiliations of 25 individuals as well as for one predicted adult mass, thus introducing a bit more uncertainty in the estimations. The fit of the model was assessed using posterior predictive checks on the predictions of individual masses (p=0.46) and survival probabilities (p=0.49). The JAGS code for this model can be found at https://github.com/timotheenivalis/SelRepSel.

## Acknowledgements

Thanks to Lauren Richardson, Tim Coulson, Alastair Wilson and Loeske Kruuk for constructive comments. Thanks to Wolf U. Blanckenhorn, Jarrod D. Hadfield, Lukas F. Keller, Marc Kéry, Hanna Kokko, Chelsea J. Little, Pirmin Nietlisbach, Barbara Tschirren and Ashley E. Latimer for comments and discussions on earlier versions of this work. Thanks also to the many field assistants. Weather data were provided by MeteoSwiss. The snow vole monitoring was authorised by the *Amt für Lebensmittelsicherheit und Tiergesundheit*, Chur.

